# Disruption of *Drosophila melanogaster* Larval Locomotion Caused by Silver Ions

**DOI:** 10.64898/2026.04.03.716380

**Authors:** Michael Stewart, Hemanta Pradhan, Xuan Zhuang, Yong Wang

**Affiliations:** Department of Physics, University of Arkansas, Fayetteville, AR 72701; Department of Biological Sciences, University of Arkansas, Fayetteville, AR 72701; Cell and Molecular Biology Program, University of Arkansas, Fayetteville, AR 72701; Materials Science and Engineering Program, University of Arkansas, Fayetteville, AR 72701

**Keywords:** locomotion, motility, fruit fly, larvae, AgNO3, AgNP

## Abstract

Silver (Ag^+^) ions are known to be toxic to bacteria, cells, organisms and living systems; yet its impacts on the locomotion of surface-crawling organisms remain poorly quantified. Here we investigated the short-term (0–6 hours) effects of Ag^+^ ions on the locomotion of *Drosophila melanogaster* larvae on flat agarose surfaces containing Ag^+^ ions at different concentrations (0, 1, 10, and 100 mM). By quantifying their locomotion, we found that *Drosophila* larvae showed shorter accumulated distances and reduced crawling speed. Additionally, we quantified the go/stop dynamics and peristalsis of the larvae and observed that Ag^+^ ions disrupted the normal, rhythmic, peristaltic contraction of the larvae and “trapped” them in the stop phase. Such toxic effects were dependent on Ag^+^ concentration and exposure duration.

## I. INTRODUCTION

The toxicity of silver (Ag) in different forms, such as Ag^+^ ions and Ag nanoparticles (AgNPs), has been extensively studied on a variety of living organisms, ranging from virus and bacteria to fungi, animal cells, microalgae, plants, and fruit flies [1–13]. It is an important topic due to the widespread use of Ag in different forms in various industries, including electronics, medicine, and consumer products [8, 14–16]. The release of Ag into the environment from these applications may pose potential risks to environmental health (particularly aquatic and terrestrial organisms), as well as human health [17, 18].

*Drosophila melanogaster*, commonly known as the fruit fly, has been used as a model organism for investigating the toxicity of Ag^+^ ions and AgNPs [1, 5, 6, 9, 10, 12, 19]. Its short life cycle (approximately 10 days from fertilization to adulthood) allows for rapid generation turnover and large-scale studies. In addition, *Drosophila* shares 60 percent of its genes with humans, many of which are associated with diseases [19, 20], enabling scientists to draw meaningful parallels between *Drosophila* and human biology. It was reported that Ag^+^ ions are more toxic than nanoparticulate silver, reducing larval survival and impairing development when administered as silver nitrate, a known ionic Ag^+^ source.[10, 12, 17] It has also been shown that AgNPs induced developmental inhibition and genotoxic effects in *Drosophila melanogaster*, affecting neural stem cells and non-neuronal cells [1, 9, 10, 12]. The AgNPs may cross the intestinal barrier, leading to increased levels of intracellular reactive oxygen species and significant DNA damage [4, 9, 10, 12, 19]. Despite these findings, the impact of Ag on the larval mobility and locomotion has been rarely investigated.

In this work, we attempted to address this knowledge gap by investigating how Ag^+^ ions affect the mobility and locomotion of *Drosophila melanogaster* larvae. We recorded the locomotion of the larvae on flat agarose surfaces containing Ag^+^ ions at different concentrations (0, 1, 10, and 100 mM) in exposure duration of 6 hours, analyzed the videos, and tracked the larvae using FIMTrack [21]. Additionally, we quantified how Ag^+^ ions changed the speeds and accumulated distances of the larvae over time. Furthermore, we analyzed the effects of Ag^+^ ions on the go/stop dynamics and peristalsis of larval body. The findings in this study showed that Ag^+^ ions reduced the locomotive ability of *Drosophila* larvae, providing new insights into Ag’s toxicity to the larval behaviors.

## II. MATERIALS AND METHODS

### A. *Drosophila* strain and maintenance

We used the *Drosophila melanogaster* strain w1118, a commonly used laboratory control background strain, originally obtained from the Bloomington Drosophila Stock Center (BDSC #5905) and maintained in the laboratory for over 5 years before use in experiments. Flies were reared on Formula 4-24 instant *Drosophila* medium (Carolina Biological Supply Co, Burlington, NC) and kept on a 12:12 h day-night cycle at a constant temperature of 26.6^°^C to ensure consistent developmental timing and physiological conditions [19, 20, 22, 23].

### B. Preparation of agarose arenas

Agarose arenas were prepared using 60 mm sterile plastic petri dishes (VWR 25384-092, Fayetteville, AR). Briefly, silver nitrate (AgNO_3_, M2AE041, Alfa Aesar) solutions were prepared fresh in deionized water, protected from light, and added to molten 1% agarose to generate final AgNO_3_ concentrations of 0 (control), 1, 10, and 100 mM. For each arena, 10 mL Ag^+^-containing 1% agarose solution was used. The prepared agarose arenas were then used within 2 hours for the behavior assays.

### C. Behavior assays

For each arena, 4–5 size-matched early third-instar feeding *Drosophila* larvae were selected and transferred to the arena immediately before the experiments. For each experiment, the arenas were placed on a circular diffusive LED disk (LUMINAIRE 09635822, Amazon) covered by a red plastic sheet (Pangda B073XJMS39, Amazon). On top of the arenas, a camera (iPhone SE 2nd Generation, Apple) was mounted on a stand, and used to record videos of the locomotion and behaviors of the *Drosophila* larvae at 0, 30, 60, 120, 180, 240, 300, and 360 min. The duration of each video was 1 min, with an exposure time of 33 ms and a frame rate of 30 fps. The magnification of the camera system corresponded to a spatial imaging scale of 16 px/mm, resulting in an effective pixel size of 0.0625 mm/px for the recorded videos. Seven replicates of the behavior experiments were performed on different days in this study. After each experiment, the *Drosophila* larvae were cooled in freezer (-20^°^C) for humane euthanasia.

### D. Data analysis

The recorded videos were analyzed using FIMTrack [21]. Raw video files were first converted into image sequences using `ffmpeg` [24]. The resultant image sequences were then imported into ImageJ, where the red channels of the images were isolated, followed by cropping the circular regions of interest (ROI, 725 px, or *∼* 48 mm, in diameter). This circular ROI corresponds to the the interior of the 60 mm petri dish. The processed image stack were saved as a TIFF file, and then loaded into FIMTrack. Tracking was performed using the standard larva model with a gray-value threshold of 40 for segmentation. The minimum and maximum larval areas were set to 125 px^2^ and 1400 px^2^, respectively, and the frame rate was set to 30 fps to match the acquisition condition. For each detected *Drosophila* larva, FIMTrack extracted the center-of-mass coordinates (*x, y*), instantaneous velocity (*v*), accumulated distance (*D*), spine length 𝓁_*s*_, and the binary go/stop phase indicator *φ*, as well as various other quantities. All these quantities from FIMTrack were used directly in further analysis. The go/stop phase indicator *φ* was used to estimate the moving ratio (*γ*) of the larvae, and the dwelling times (*τ*_go_ and *τ*_stop_). Additionally, the trajectories of spine length 𝓁_*s*_ were subjected to Hilbert transform [25–27] for estimating the amplitudes (*A*_*s*_) and frequencies (*f*_*s*_) of the oscillation of *Drosophila* larvae’s locomotion.

## III. RESULTS

### A. Trajectories of *Drosophila* larvae on Ag^+^-containing agarose surface

Several examples of the crawling trajectories of *Drosophila* larvae on the agarose arenas containing different concentrations of Ag^+^ ions are shown in Fig. 2 and S1, where each colored curve represents the trajectory of a single larva. The controls (i.e., in the absence of Ag^+^ ions, or 0 mM) showed long, continuous trajectories that traversed large portions of the arenas (Fig. 2). Most of the larvae displayed sustained crawling activity: some larvae exhibited smooth extended paths, while others showed frequent directional changes (Fig. 2). During the 6-hour period, both the path lengths and the fraction of larvae showing long paths decreased; however, the overall exploratory behavior remained robust (Fig. 2). In contrast, larvae exposed to Ag^+^ ions displayed shorter trajectories and reduced mobility. The reduction was both dose-dependent and time-dependent. In the presence of 1 mM Ag^+^ ions, the crawling paths of the larvae were shorter than the controls but many larvae continued to explore the arenas, with trajectories gradually contracting over time (Fig. 2). In the presence of 10 mM Ag^+^ ions, the movement of the larvae became markedly re-stricted, producing short, fragmented segments at the proximity of their starting locations (Fig. 2). The crawling paths of the larvae further diminished as the exposure time increased. Lastly, in the presence of 100 mM Ag^+^ ions, the larvae showed very brief initial movement before becoming completely immobile, resulting in collapsed trajectories (Fig. 2).

**FIG. 1.**
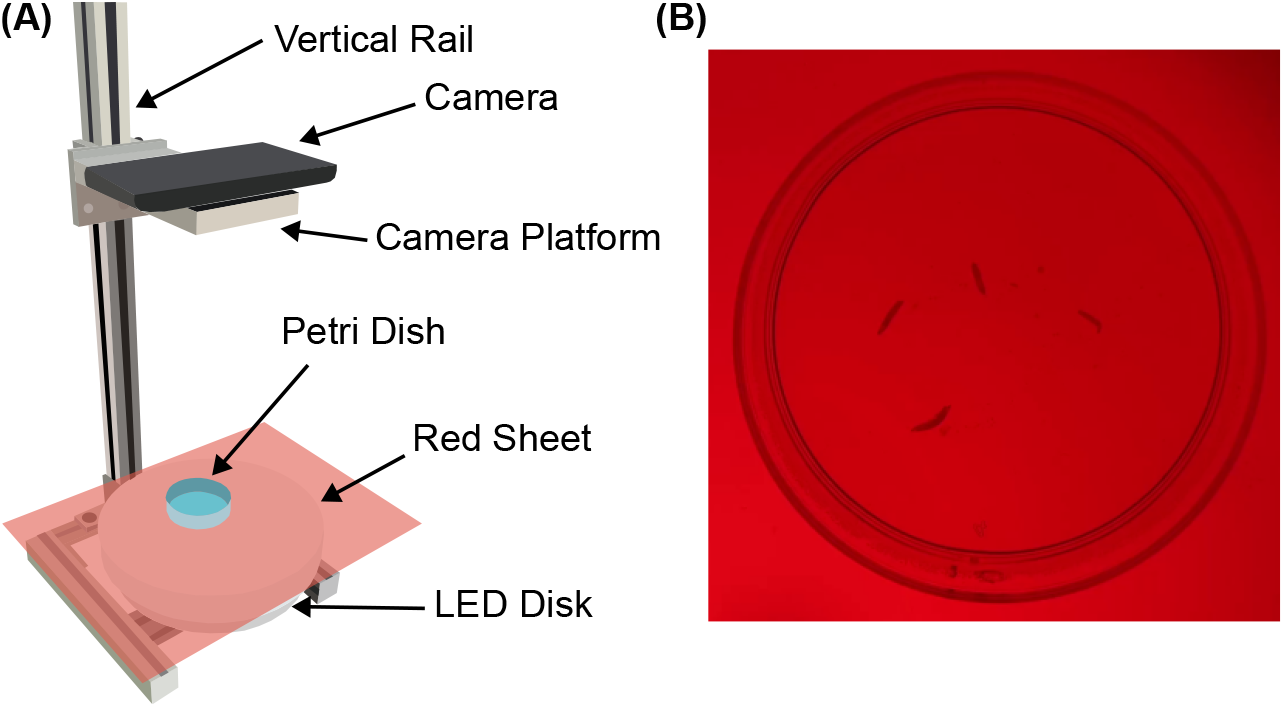
Illustration of imaging *Drosophila* larvae on Ag^+^-containing agarose arenas. **(A)** Schematic diagram of the imaging apparatus. **(B)** Examples of raw images of the *Drosophila* larvae on agarose arenas containing AgNO_3_ at different concentrations.

**FIG. 2.**
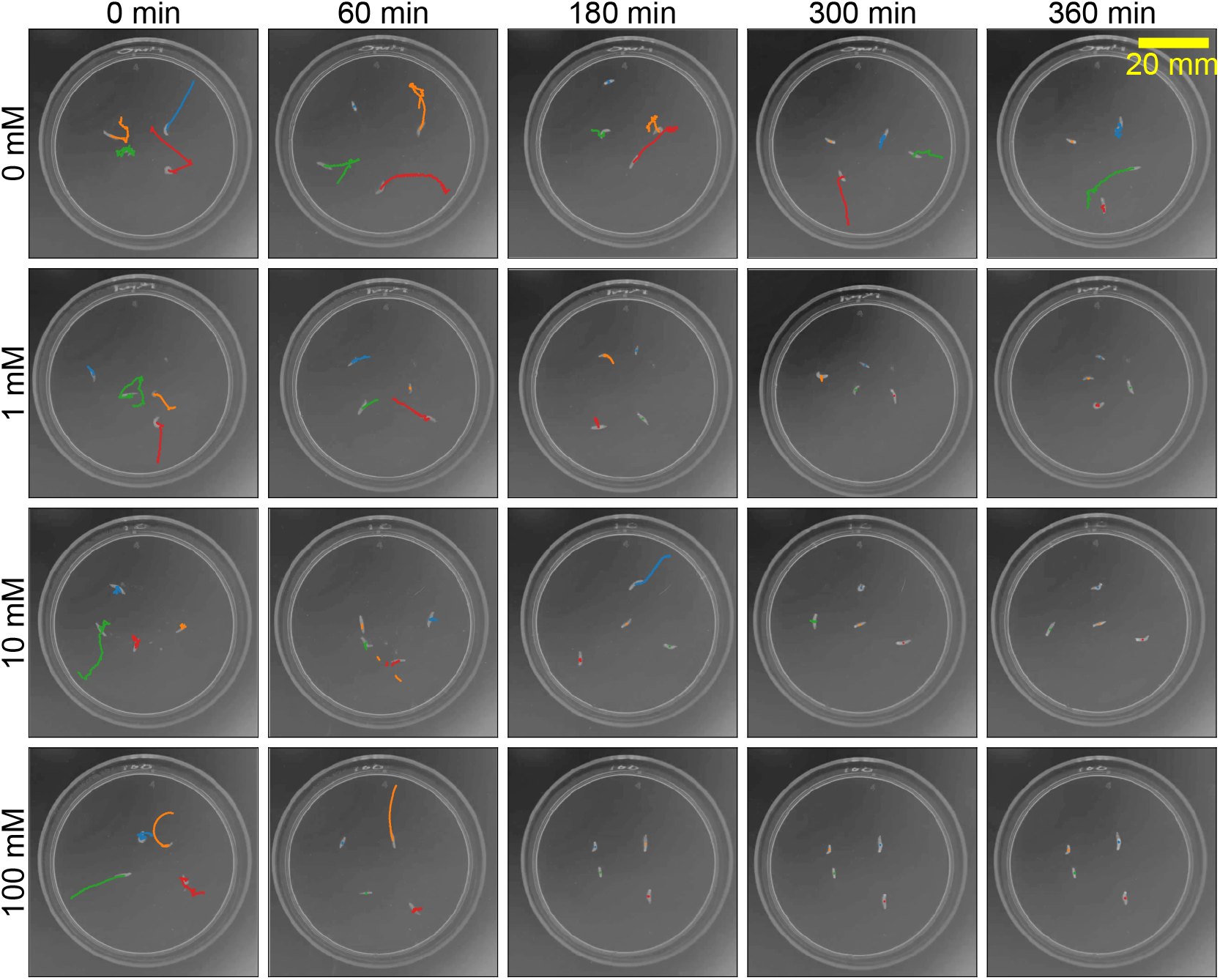
Examples of trajectories of *Drosophila* larvae (colored curves) in the absence (0 mM) and presence (1 mM, 10 mM, and 100 mM) of Ag^+^ ions for different durations (0 min, 60 min, 180 min, 300 min, and 360 min). Scale bar = 20 mm.

### B. Ag^+^ ions shortened the accumulated distances crawled by *Drosophila* larvae

To quantify the observations from the behavior assays (Fig. 2), we examined the accumulated distances over one minute (*D*) of the *Drosophila* larvae. As shown in Fig. 3A, the accumulated distances (*D*) of the *Drosophila* larvae in the absence of Ag^+^ ions declined only slightly. However, the larvae exposed to Ag^+^ ions at 1 mM, 10 mM, and 100 mM exhibited markedly steeper reductions over time. After 6 hours, the accumulated distances over one minute (*D*) of the larvae exposed to Ag^+^ ions were less than 50% of that of the controls (Fig. 3B). The changes in the accumulated distances over one minute during the 6-hour experimental period (Δ*D* = *D*_360_ − *D*_0_) confirmed that the reduction in accumulated distance over one minute was much more significant for *Drosophila* larvae exposed to Ag^+^ ions than the controls in the absence of Ag^+^ ions (Fig. 3C).

**FIG. 3.**
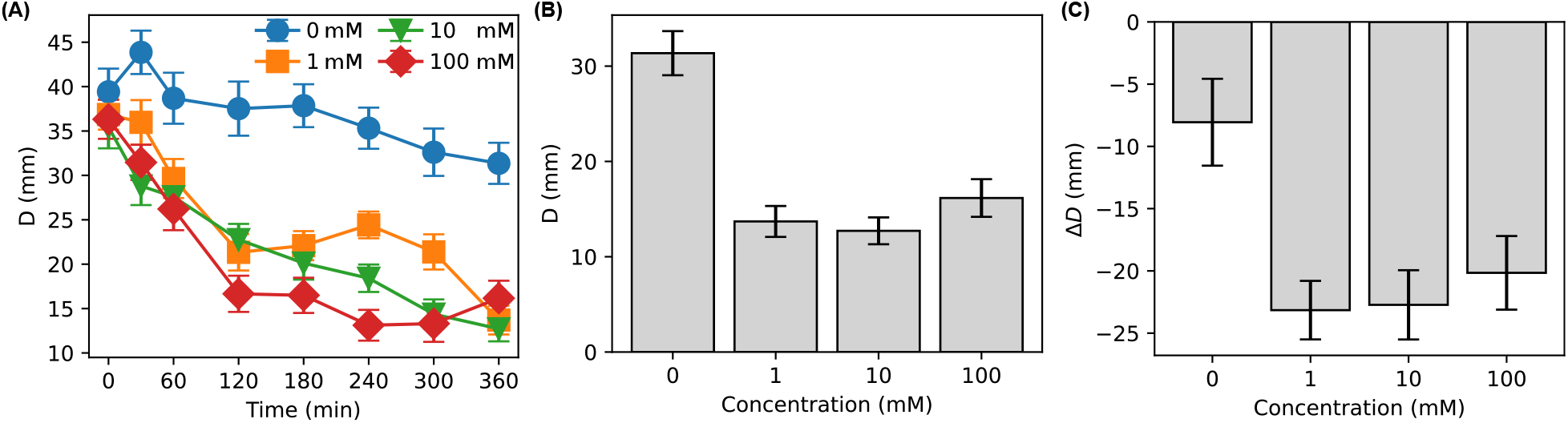
**(A)** Average accumulated distance *D* of *Drosophila* larvae in one minute as functions of time in the presence of Ag^+^ ions of different concentrations (0 – 100 mM). **(B)** Comparison of accumulated distance *D* of *Drosophila* larvae in one minute after exposure to Ag^+^ ions of different concentrations (0 – 100 mM) for 6 hours. **(C)** Changes of accumulated distance Δ*D* of *Drosophila* larvae in one minute after exposure to Ag^+^ ions of different concentrations (0 – 100 mM) for 6 hours. Error bars in all panels indicate standard errors of the means (SEM).

### C. Ag^+^ ions reduced the speed of *Drosophila* larvae

We also examined the effect of Ag^+^ ions on the speed of *Drosophila* larvae. Similar to the accumulated distance, the average speed (*v*) of larvae in the absence of Ag^+^ ions (0 mM, the controls) remained relatively stable over the 6-hour experimental period (Fig. 4A). In contrast, the larvae exposed to Ag^+^ ions at 1 mM, 10 mM, and 100 mM showed reduced speeds (Fig. 4A). At the end of 6-hour experimental period, the average speed of larvae exposed to Ag^+^ ions was less than 25% of that of the controls (Fig. 4B).

**FIG. 4.**
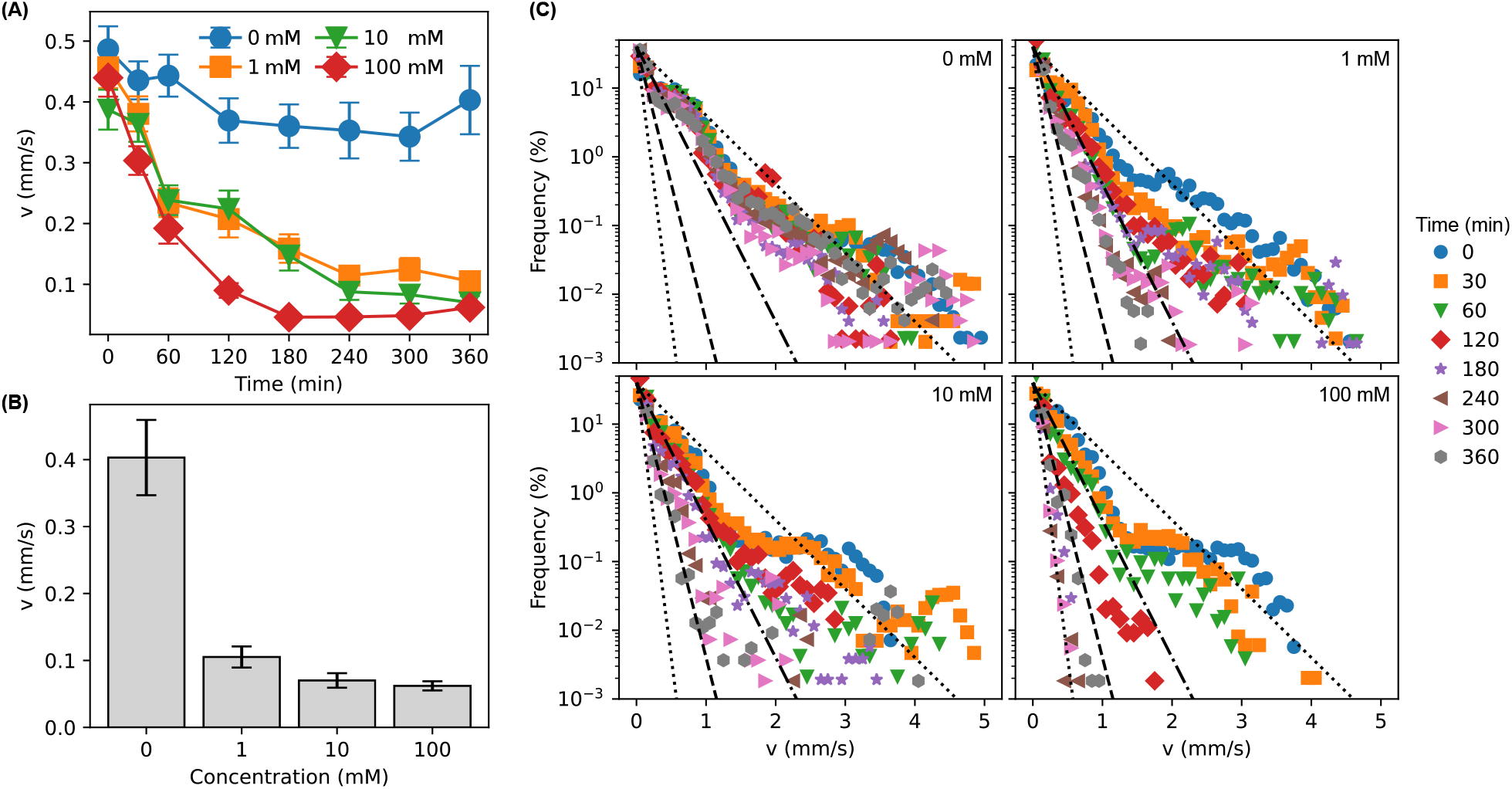
**(A)** Average speed *v* of *Drosophila* larvae as functions of time in the presence of Ag^+^ ions of different concentrations (0 – 100 mM). **(B)** Comparison of *Drosophila* larvae speed after exposure to Ag^+^ ions of different concentrations (0 – 100 mM) for 6 hours. Error bars in panels A and B indicate SEM. **(C)** Distributions of *Drosophila* larvae speed in the absence (0 mM) and presence (1 mM, 10 mM, and 100 mM) of Ag^+^ ions for different durations (0 – 360 min) in log-linear scale. The dotted, dashed, dashed-dotted lines in panel C indicate slopes of −1, −2, −4, and −8, in the log-linear scale.

In addition to the average larvae speeds, we investigated the distribution of instantaneous speed of the *Drosophila* larvae. As shown in Fig. 4C, the distributions of the instantaneous speed of larvae without exposure to Ag^+^ ions followed roughly exponential decays (*∼ e*^−*kv*^), and the slope *k* in the log-linear scale was between −1 and −2 (Fig. 4C). In addition, these distributions in the control did not change significantly over the 6-hour experimental period (Fig. 4C). In contrast, exposure to Ag^+^ ions led to faster decays in the instantaneous speed distributions (Fig. 4C) as the exposure time increased. For example, the slope in the log-linear scale for the larvae exposed to 10 mM Ag^+^ ions reached −4 at 6 hours (Fig. 4C). Steeper distributions of the larvae instantaneous speed were consistent with the reduction in the average speed (Fig. 4A and B). Another observation is that the instantaneous speed distributions of the larvae were affected by Ag^+^ ions as soon as they were placed on the Ag^+^-containing agarose arenas (*t* = 0 min, Fig. 4C), especially with Ag^+^ ions at 10 mM and 100 mM. This short time scale suggests that the early behavioral effect of Ag^+^ ions on *Drosophila* larvae may involve acute sensory or contact-dependent responses.

### D. Ag^+^ ions disrupt go/stop dynamics of *Drosophila* larvae

To obtain a deeper understanding of the effects of Ag^+^ ions on the locomotion of *Drosophila* larvae, we analyzed their go/stop dynamics in detail, based on the go/stop phases obtained from the FIMTrack program [21], in which the go phase of a larva was identified if its mean velocity (*v*_*m*_) was above 6 pixels per 10 frames and its bending angle is below 30^°^. An example of the go/stop phase (*φ*) trajectory is shown in Fig. 5A, together with the associated trajectory of the instantaneous speed. From the go/stop phase trajectory, we calculated the moving ratio, *γ* = *N*_go_*/*(*N*_go_ + *N*_stop_), where *N*_go_ and *N*_stop_ are the numbers of states in the go and stop phases, respectively. We observed that the average moving ratio of *Drosophila* larvae in the absence of Ag^+^ ions decreased only slightly (Fig. 5B). In contrast, the moving ratio of larvae exposed to Ag^+^ ions declined dramatically to *∼* 0% in 6 hours (Fig. 5B). With 100 mM Ag^+^ ions, larvae remained in the stop phase completely at 3 hours. The trends observed for the moving ratio were consistent with those for the accumulated distances and speeds of the larvae (Fig. 3 and 4).

**FIG. 5.**
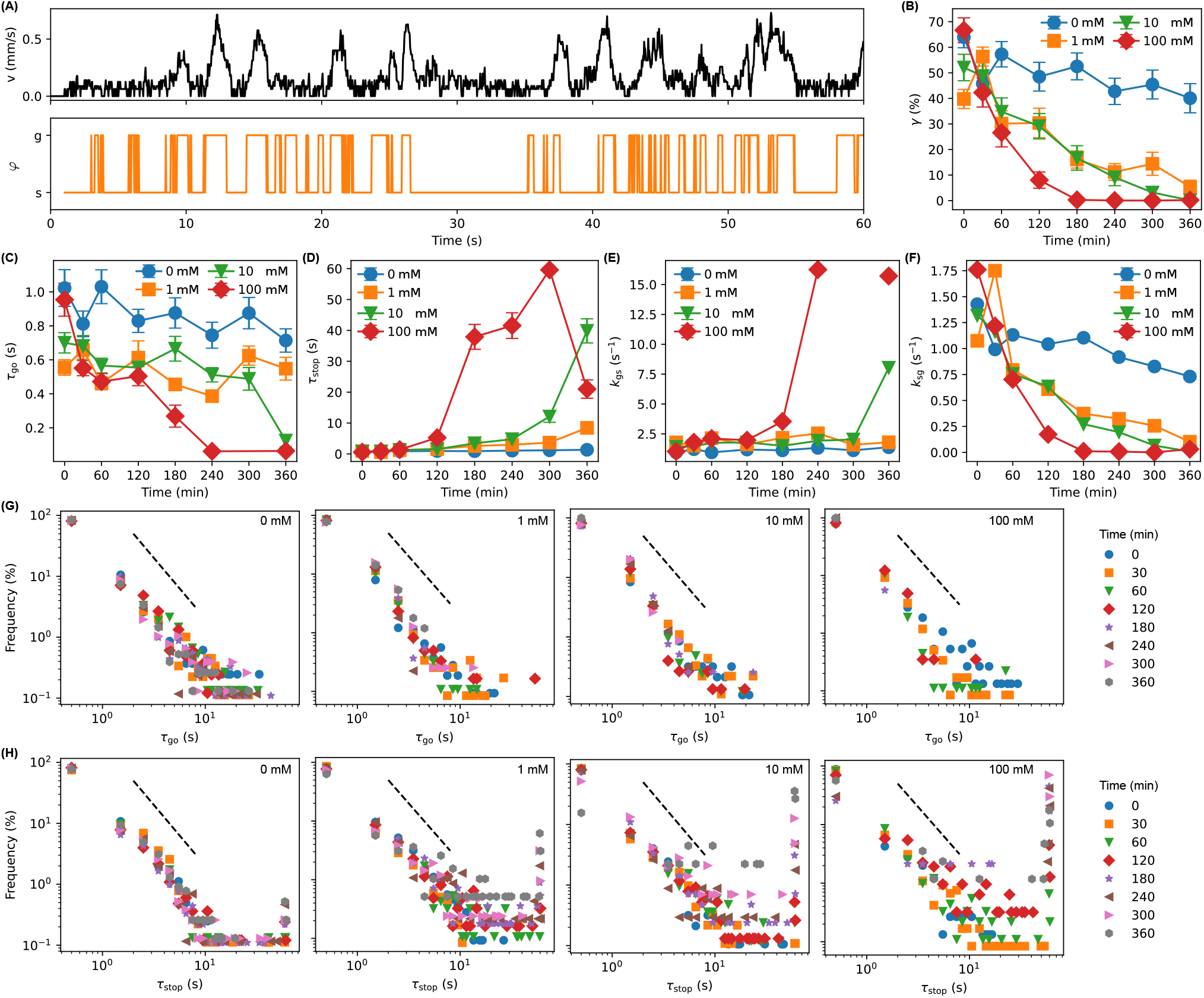
**(A)** An example of the speed and go-phase trajectory of a *Drosophila* larva, obtained from FIMTrack [21]. **(B)** Average go-phase ratios *γ* of *Drosophila* larvae as functions of time in the presence of Ag^+^ ions of different concentrations (0 – 100 mM). **(C, D)** Average dwelling times in the (C) go and (D) stop phases (*τ*_go_ and *τ*_stop_, respectively) of *Drosophila* larvae as functions of time in the presence of Ag^+^ ions of different concentrations (0 – 100 mM). (**E, F**) Estimated transition rates (*k*_*gs*_ and *k*_*sg*_) of *Drosophila* larvae as functions of time in the presence of Ag^+^ ions of different concentrations. Error bars in panels B–F indicate SEM. **(G, H)** Distributions of *Drosophila* larvae dwelling times (*τ*_go_ and *τ*_stop_) in the absence (0 mM) and presence (1 mM, 10 mM, and 100 mM) of Ag^+^ ions for different durations (0 – 360 min) in log-log scale. The dashed lines in panels G and H indicate slopes of −2 in the log-log scale.

Additionally, we determined the dwelling times in the go and stop phases (*τ*_go_ and *τ*_stop_) from the go/stop phase trajectories. We observed that the average dwelling times of the larvae in the absence of Ag^+^ ions, or exposed to 1 mM Ag^+^ ions, remained roughly constant (blue circles and orange squares in Fig. 5C and D). In contrast, the effects of Ag^+^ ions at 10 mM and 100 mM were much more obvious: *τ*_go_ decreased and *τ*_stop_ increased as the exposure time increased (green triangles and red diamonds in Fig. 5C and D).

Furthermore, we estimated the transition rates (*k*_*gs*_ and *k*_*sg*_) between the go and stop phases from the phase trajectories by *k*_*gs*_ = *P*_*gs*_*/*Δ*t* and *k*_*sg*_ = *P*_*sg*_*/*Δ*t*, where *P*_*gs*_ and *P*_*sg*_ were the probabilities of the larvae from go phase to stop phase or vice versa, respectively, and Δ*t* was the time interval between frames. As shown in Fig. 5E and F, we found that the go-to-stop transition rate increased significantly for larvae exposed to 10 and 100 mM Ag^+^ ions for 6 hours, while it remained roughly constant for untreated larvae and larvae exposed to 1 mM Ag^+^ ions. Interestingly, even at 1 mM Ag^+^ ions, the stop-to-go transition rate of the larvae dramatically decreased to close to 0 s^−1^ after 6 hours. This finding indicates that Ag^+^ ions “trapped” the larvae in the stop phase, reducing the speed and accumulated distances as observed in Fig. 3 and 4.

Lastly, we examined the distributions of the dwelling times in Fig. 5G and H, which showed power law distributions with a power of roughly −2 (dashed line in Fig. 5G and H). Similar power law distributions have been commonly observed in living systems, reflecting their complex, non-Markov nature [28–30]. Interestingly, Ag^+^ ions did not change the exponent in the power laws, although they led to much heavier tails for the stop phase dwelling time (Fig. 5G and H), suggesting that the larvae were “trapped” in the stop phase after exposure to Ag^+^ ions.

### E. Ag^+^ ions interrupt the peristalsis of *Drosophila* larvae

Due to the importance of peristalsis (the rhythmic wave-like contraction) of *Drosophila* larvae for their movement and survival [30–32], we quantified and compared the peristaltic wave dynamics of the larvae after exposure to Ag^+^ ions, based on the measured spine lengths (𝓁_*s*_) of the larvae. An example of larvae spine-length trajectories is shown in Fig. 6A (blue solid line), which exhibits noisy oscillations with gradual shifts. Before estimating the amplitude and frequency of the oscillations, we calculated the baseline (red dashed line in Fig. 6A) by applying a median filter and removed the baseline from the trajectory, which resulted in a detrended trajectory of the spine-length (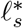, black solid line in Fig. 6A). Then, the Hilbert transform [25–27] was applied to the detrended trajectory and the instantaneous amplitudes (*A*_*s*_, orange solid line in Fig. 6A) and instantaneous frequencies (*f*_*s*_, green solid line in Fig. 6A) were obtained. The average value of the instantaneous frequencies (red dashed line in Fig. 6B) was used to report the frequency of the peristalsis of the larva. Note that the mean value was consistent with both the center peak of the distribution of the instantaneous frequencies, and the manual measurement from the original trajectory of the larval spine length.

**FIG. 6.**
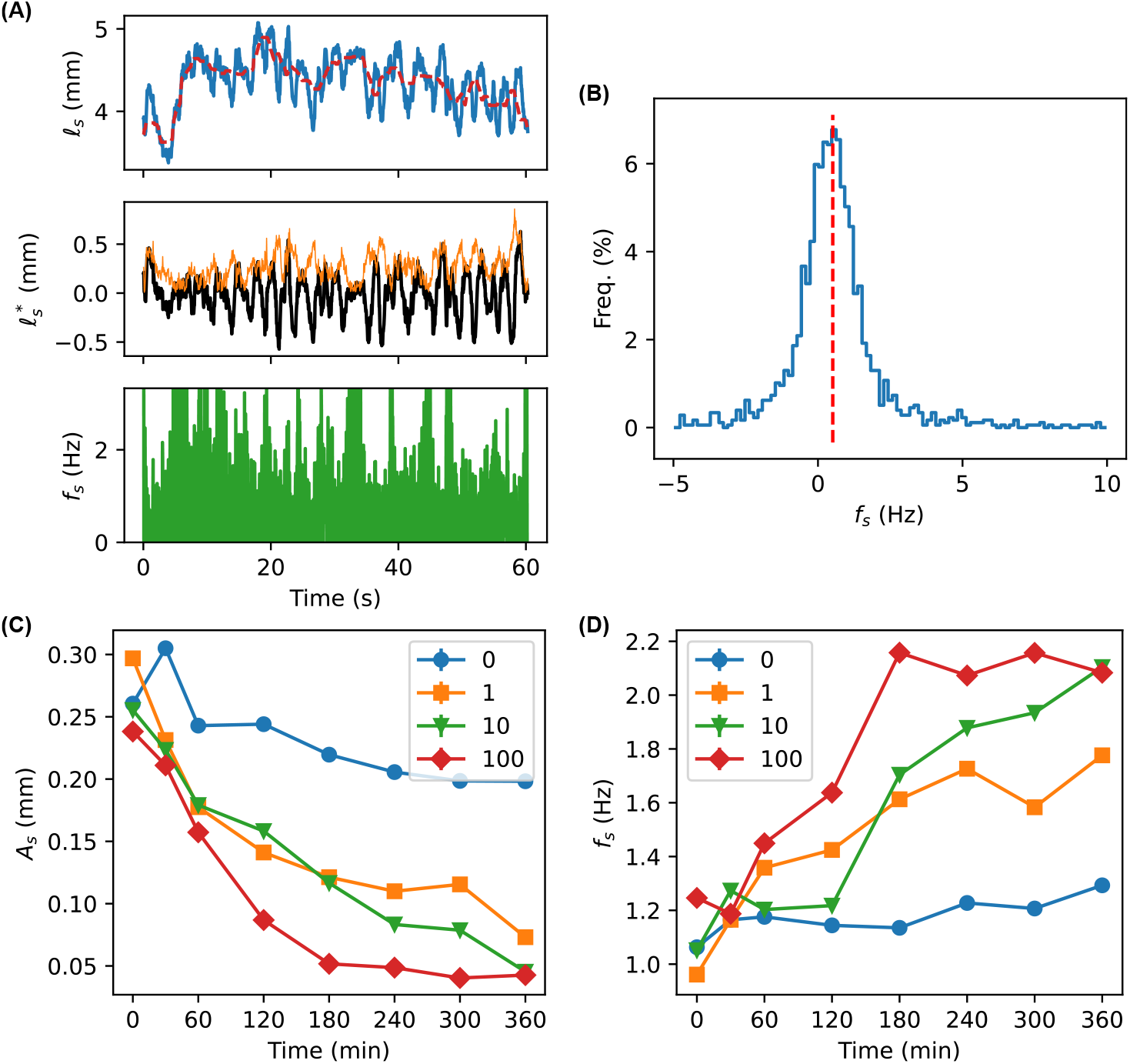
**(A)** An example of the spine length trajectory of a *Drosophila* larva, before (blue curve, 𝓁_*s*_) and after (black curve, 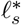) applying median filter detrending (red dashed curve indicating the baseline for detrending), and the instantaneous amplitudes (orange curve, *A*_*s*_) and frequencies (green curve, *f*_*s*_) calculated from Hilbert transform. **(B)** Distribution of the instantaneous frequency *f*_*s*_ from panel A. The red dashed line indicate the average of 0.5 Hz. **(C, D)** Estimated averages of (C) amplitudes and (D) frequencies of *Drosophila* larvae spine lengths as functions of time in the presence of Ag^+^ ions of different concentrations. Error bars (smaller than symbols) in panels C and D indicate SEM.

We observed that the mean oscillation amplitude decreased as the exposure time of larvae to Ag^+^ ions increased (Fig. 6C). Although the larvae without Ag^+^ ions showed slight decrease in the oscillation amplitude, the changes in the Ag^+^-treated larvae were much more significant (Fig. 6C). In addition to the reduced oscillation amplitude, we observed that exposure to Ag^+^ ions led to an apparent increase in oscillation frequency of the spine-length (Fig. 6D).

## IV. CONCLUSIONS AND DISCUSSIONS

To summarize, we investigated the acute effects of Ag^+^ ions on the locomotion of *Drosophila* larvae based on a behavior assay on Ag^+^-containing agarose arenas and quantitative analysis. Upon exposure to Ag^+^ ions, *Drosophila* larvae showed shorter accumulated distances (Fig. 3) and reduced crawling speed (Fig. 4). More importantly, the go/stop dynamics (Fig. 5) and peristalsis (Fig. 6) of the larvae were disrupted by the Ag^+^ ions. Additionally, dose dependence and time dependence were observed for these effects.

*Drosophila melanogaster* larvae struggled to generate the normal, rhythmic, peristaltic locomotion when crawling on Ag^+^-containing agarose surfaces. We observed that Ag^+^ ions caused decreases in the amplitude, and increases in the frequency, of spine-length oscillations. Because spine-length oscillation is a proxy for peristaltic wave dynamics, the observed changes provided evidence of disrupted locomotor rhythm, and possibly alterations in the muscle contraction dynamics [30–34]. These disruptions were likely to reduce and diminish the efficiency of the larvae to generate forward locomotion, explaining the observed decreases in the moving ratio and the go phase dwelling time, elongation of the stop phase dwelling time, and reductions in the speed and accumulated distances of the larvae after exposure to Ag^+^ ions. It would be interesting to examine the effect of Ag^+^ ions on the muscle contraction dynamics of larvae directly in future studies. Additionally, as differences were observed between larvae on Ag^+^-absent and Ag^+^-containing agarose arenas as soon as the larvae were transferred to the arenas, the changes in their locomotion may involve acute, contact-dependent sensory responses, including gustatory, nociceptive, or other peripheral sensory pathways, besides the impairment developed during continued exposure. It would be interesting to examine the mechanism of Ag^+^’s effects on these sensory pathways in the future.

Ag^+^ ions are known to be toxic to various biological systems [16, 17, 35, 36] by reducing ATP production [4, 7, 16, 36], damaging cell membrane and DNA [3, 10, 12, 16, 36], inducing oxidative and metabolic stresses [4, 9, 10, 16, 36], disrupting calcium handling [4, 7, 16, 36], and interfering with neuromuscular junction signaling [10, 32, 34, 36]. It would be exciting to investigate the mechanism for the observed disruption of larval locomotion upon exposure to Ag^+^ ions in the future. For example, combining calcium imaging, electrophysiology, and genetic perturbations is expected to facilitate mechanistic studies on the Ag^+^-induced disruption of larval locomotion. It would also be interesting to compare how and why different heavy metal ions (e.g., Ag^+^, Cu^2+^, Au^+^, Zn^2+^) affect the locomotion of *Drosophila* larvae.

The concentrations Ag^+^ ions used in the current study were high in order to observe their short-term, acute effects on the locomotion of *Drosophila* larvae. It would be interesting to examine in the future the long-term effects of Ag^+^ ions at lower concentrations. Additionally, as the toxicity of silver nanoparticles (AgNPs) is partly due to the released Ag^+^ ions [37, 38], we expected that *Drosophila* larvae exposed to AgNPs would show similar changes in locomotion observed in this study. However, it has been reported that AgNPs show particle-specific toxicity to bacteria compared to Ag^+^ ions [2, 3, 8, 39]; thus it is worthwhile studying how and why AgNPs interfere with larval locomotion.

Several potential practical applications are expected from the current study. For example, silver-based pesticide and technology may be developed and optimized to target harmful insects to reduce their larval mobility and survival to enhance crop protection in agriculture, or to reduce disease transmission in pest control [40, 41].

## Acknowledgment

This work was not supported directly by research grants. We are grateful for support from the Arkansas High Performance Computing Center (AHPCC), which is funded in part by the National Science Foundation and the Arkansas Science and Technology Authority. XZ was supported by the National Institutes of Health (grant No. R15GM152956 and R35GM160135).

## Author Contributions

MS and YW conceived and designed the project; MS performed experiments and acquired the data; MS, HP, and YW performed data analysis and visualization; MS and YW wrote the paper; all authors reviewed, commented, and revised the paper.

## SUPPLEMENTARY INFORMATION

**FIG. S1.**
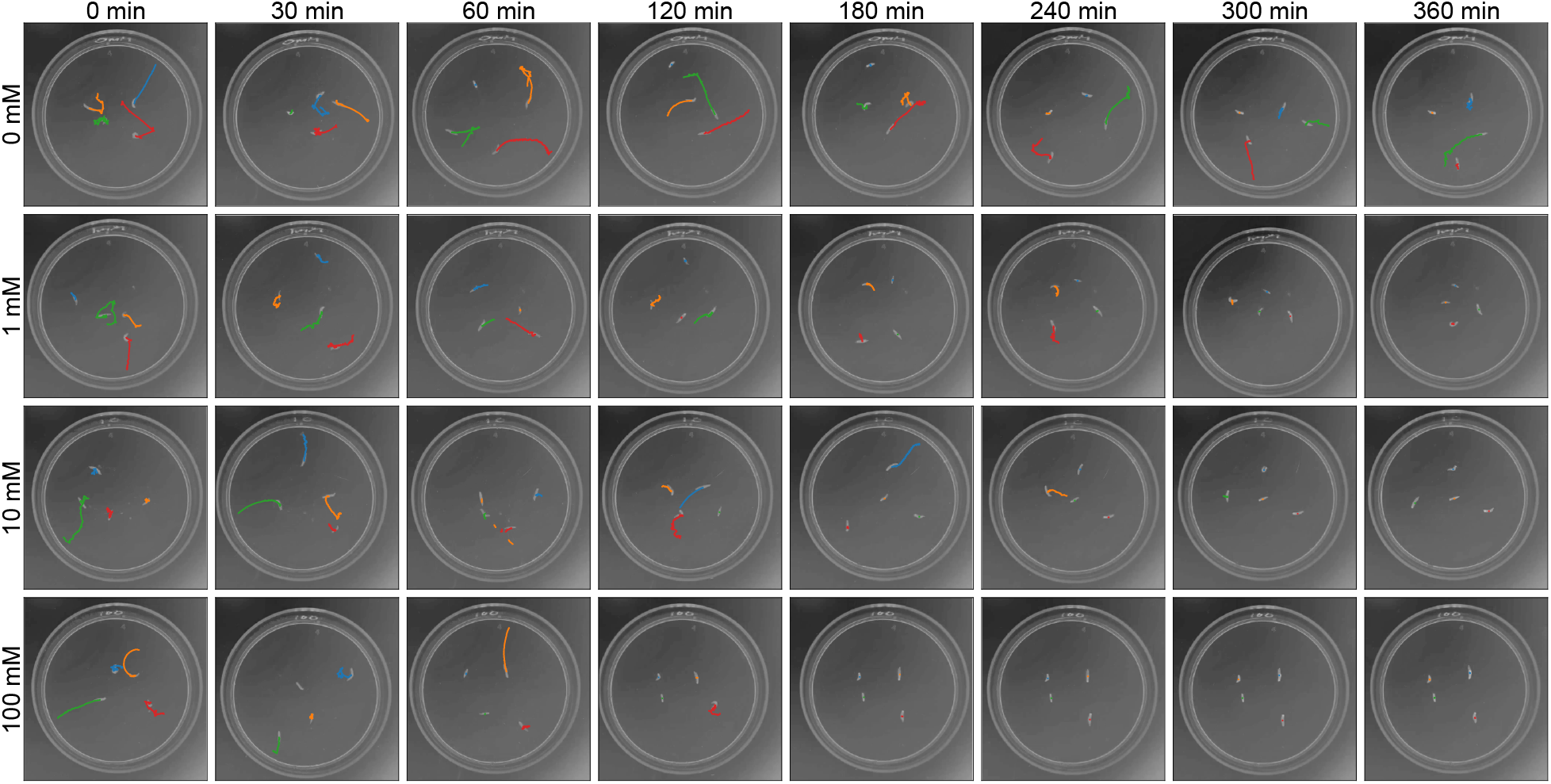
Examples of trajectories of Drosophila larvae (colored curves) in the absence (0 mM) and presence (1 mM, 10 mM, and 100 mM) of Ag^+^ ions for different durations (0 min, 60 min, 120 min, 180 min, 240 min, 300 min, and 360 min).

## References

[1] A. Panacek, R. Prucek, D. Safarova, M. Dittrich, J. Richtrova, K. Benickova, R. Zboril, and L. Kvitek, Acute and chronic toxicity effects of silver nanoparticles (nps) on drosophila melanogaster, Environmental Science & Technology 45, 4974 (2011).

[2] Y. Dong, H. Zhu, Y. Shen, W. Zhang, and L. Zhang, Antibacterial activity of silver nanoparticles of different particle size against vibrio natriegens, PLOS ONE 14, e0222322 (2019).

[3] A. D. Burchardt, R. N. Carvalho, A. Valente, P. Nativo, D. Gilliland, C. P. García, R. Passarella, V. Pedroni, F. Rossi, and T. Lettieri, Effects of silver nanoparticles in diatom Thalassiosira pseudonana and cyanobacterium Synechococcus sp., Environmental Science & Technology 46, 11336 (2012).

[4] Y. Huang, R. Chen, Y. Chen, and X. Lü, Investigation of molecular mechanisms in silver nanoparticle-induced cytotoxicity from gene to metabolite level, Scientific Reports 15, 26923 (2025).

[5] Z. Wang, L. Zhang, and X. Wang, Molecular toxicity and defense mechanisms induced by silver nanoparticles in Drosophila melanogaster, Journal of Environmental Sciences 125, 616 (2023).

[6] D. J. Gorth, D. M. Rand, and T. J. Webster, Silver nanoparticle toxicity in drosophila: size does matter, International Journal of Nanomedicine 6, 343 (2011).

[7] E.-J. Park, J. Yi, Y. Kim, K. Choi, and K. Park, Silver nanoparticles induce cytotoxicity by a trojan-horse type mechanism, Toxicology in Vitro 24, 872 (2010).

[8] P. R. More, S. Pandit, A. De Filippis, G. Franci, I. Mijakovic, and M. Galdiero, Silver nanoparticles: Bactericidal and mechanistic approach against drug resistant pathogens, Microorganisms 11, 10.3390/microorganisms11020369 (2023).

[9] M. Jalal, S. H. Haidar, S. Rafique, S. Azan, F. Batool, and N. Mehmood, The effect of silver nanoparticles on oxidative stress enzymes in Drosophila melanogaster, Scholars Academic Journal of Biosciences 13, 614 (2025).

[10] M. Alaraby, S. Romero, A. Hernández, and R. Marcos, Toxic and genotoxic effects of silver nanoparticles in Drosophila, Environmental and Molecular Mutagenesis 60, 277 (2019).

[11] G. Waktole, Toxicity and molecular mechanisms of actions of silver nanoparticles, Journal of Biomaterials and Nanobiotechnology 14, 53 (2023).

[12] S.-E. A. Araj, N. M. Salem, I. H. Ghabeish, and A. M. Awwad, Toxicity of nanoparticles against Drosophila melanogaster (diptera: Drosophilidae), Journal of Nanomaterials 2015, 758132 (2015).

[13] M. J. Akhtar, M. Ahamed, and H. A. Alhadlaq, Understanding the silver nanotoxicity: Mechanisms, risks, and mitigation strategies, Journal of Nanoparticle Research 27, 1 (2025).

[14] H. Y. Lee, H. K. Park, Y. M. Lee, K. Kim, and S. B. Park, A practical procedure for producing silver nanocoated fabric and its antibacterial evaluation for biomedical applications, Chemical Communications (Cambridge, England), 2959 (2007), 17622444.

[15] N. Vigneshwaran, A. A. Kathe, P. V. Varadarajan, R. P. Nachane, and R. H. Balasubramanya, Functional finishing of cotton fabrics using silver nanoparticles, Journal of Nanoscience and Nanotechnology 7, 1893 (2007).

[16] X. Chen and H. J. Schluesener, Nanosilver: A nanoproduct in medical application, Toxicology Letters 176, 1 (2008).

[17] B. K. Gaiser, T. F. Fernandes, M. Jepson, J. R. Lead, C. R. Tyler, and V. Stone, Assessing exposure, uptake and toxicity of silver and cerium dioxide nanoparticles from contaminated environments, Environmental Health 8, S2 (2009).

[18] D. Samal, P. Khandayataray, M. Sravani, and M. K. Murthy, Silver nanoparticle ecotoxicity and phytoremediation: A critical review of current research and future prospects, Environmental Science and Pollution Research 31, 8400 (2024).

[19] C. Ong, L.-Y. L. Yung, C. Yu, B.-H. Bay, and G.-H. Baeg, Drosophila melanogaster as a model organism to study nanotoxicity, Nanotoxicology 9, 396 (2015).

[20] U. B. Pandey and C. D. Nichols, Human disease models in Drosophila melanogaster and the role of the fly in therapeutic drug discovery, Pharmacological Reviews 63, 411 (2011), 21415126.

[21] B. Risse, D. Berh, N. Otto, C. Klämbt, and X. Jiang, FIMTrack: An open source tracking and locomotion analysis software for small animals, PLOS Computational Biology 13, e1005530 (2017).

[22] H. J. MacLean, J. Overgaard, T. N. Kristensen, C. Lyster, L. Hessner, E. Olsvig, and J. G. Sørensen, Temperature preference across life stages and acclimation temperatures investigated in four species of Drosophila, Journal of Thermal Biology 86, 102428 (2019).

[23] F. Missirlis, Regulation and biological function of metal ions in drosophila, Current Opinion in Insect Science Global Change Biology. Molecular Physiology. October (2021), 47, 18 (2021).

[24] S. Tomar, Converting video formats with ffmpeg, Linux Journal 2006, 10 (2006).

[25] D. Benitez, P. A. Gaydecki, A. Zaidi, and A. P. Fitzpatrick, The use of the hilbert transform in ecg signal analysis, Computers in Biology and Medicine 31, 399 (2001).

[26] Y.-W. Liu, Hilbert transform and applications, in Fourier Transform - Applications, edited by S. M. Salih (IntechOpen, London, 2012) Chap. 12.

[27] V. Gogula and B. Edward, Advanced signal analysis for high-impedance fault detection in distribution systems: a dynamic hilbert transform method, Frontiers in Energy Research Volume 12 - 2024, 10.3389/fenrg.2024.1365538 (2024).

[28] A. Clauset, C. R. Shalizi, and M. E. J. Newman, Power-law distributions in empirical data, SIAM Review 51, 661 (2009).

[29] J. M. Moore, G. Yan, and E. G. Altmann, Nonparametric power-law surrogates, Physical Review X 12, 021056 (2022).

[30] J. Loveless, K. Lagogiannis, and B. Webb, Modelling the mechanics of exploration in larval Drosophila, PLOS Computational Biology 15, e1006635 (2019).

[31] J. C. Caldwell, M. M. Miller, S. Wing, D. R. Soll, and D. F. Eberl, Dynamic analysis of larval locomotion in Drosophila chordotonal organ mutants, Proceedings of the National Academy of Sciences 100, 16053 (2003).

[32] M. Q. Clark, A. A. Zarin, A. Carreira-Rosario, and C. Q. Doe, Neural circuits driving larval locomotion in Drosophila, Neural Development 13, 6 (2018).

[33] M. N. Günther, G. Nettesheim, and G. T. Shubeita, Quantifying and predicting Drosophila larvae crawling phenotypes, Scientific Reports 6, 27972 (2016).

[34] Y. Guo, Y. Wang, W. Zhang, S. Meltzer, D. Zanini, Y. Yu, J. Li, T. Cheng, Z. Guo, Q. Wang, J. S. Jacobs, Y. Sharma, D. F. Eberl, M. C. Göpfert, L. Y. Jan, Y. N. Jan, and Z. Wang, Transmembrane channel-like (tmc) gene regulates Drosophila larval locomotion, Proceedings of the National Academy of Sciences of the United States of America 113, 7243 (2016).

[35] E.-J. Park, S. J. Lee, G.-H. Lee, D.-W. Kim, C. Yoon, B.-S. Lee, Y. Kim, J. Chang, and K. Lee, Comparison of subchronic immunotoxicity of four different types of aluminum-based nanoparticles, Journal of Applied Toxicology 38, 575 (2018).

[36] B. Reidy, A. Haase, A. Luch, K. A. Dawson, and I. Lynch, Mechanisms of silver nanoparticle release, transformation and toxicity: A critical review of current knowledge and recommendations for future studies and applications, Materials 6, 2295 (2013).

[37] Y. Lee, J. Kim, J. Oh, S. Bae, S. Lee, I. S. Hong, and S. Kim, Ion-release kinetics and eco-toxicity effects of silver nanoparticles*, Environmental Toxicology and Chemistry 31, 155 (2011), https://academic.oup.com/etc/article-pdf/31/1/155/61001022/717.pdf.

[38] S. Kittler, C. Greulich, J. Diendorf, M. Köller, and M. Epple, Toxicity of silver nanoparticles increases during storage because of slow dissolution under release of silver ions, Chemistry of Materials 22, 4548 (2010).

[39] Z.-m. Xiu, Q.-b. Zhang, H. L. Puppala, V. L. Colvin, and P. J. J. Alvarez, Negligible particle-specific antibacterial activity of silver nanoparticles, Nano Letters 12, 4271 (2012).

[40] C. J. Osborne, A. E. Norton, R. J. Whitworth, K. S. Silver, and L. W. Cohnstaedt, Tiny silver bullets: silver nanoparticles are insecticidal to Culicoides sonorensis (diptera: Ceratopogonidae) biting midge larvae, Journal of Medical Entomology 61, 1427 (2024).

[41] D. Martínez-Cisterna and colleagues, Silver nanoparticles as a potent nanopesticide: Toxic effects and action mechanisms on pest insects of agricultural importance—a review, Molecules 29, 5520 (2024).

